# Kinetic and thermodynamic allostery in the Ras protein family

**DOI:** 10.1101/2022.11.29.518294

**Authors:** Leigh J. Manley, Milo M. Lin

## Abstract

Allostery, the tranfer of information between distant parts of a macromolecule, is a fundamental feature of protein function and regulation. However, allosteric mechanisms are usually not explained by protein structure, requiring information on correlated fluctuations uniquely accessible to molecular simulation. Existing work to extract allosteric pathways from molecular dynamics simulations has focused on thermodynamic correlations. Here we show how kinetic correlations (i.e. dynamical activity) encode complementary information essential to explain observed variations in allosteric regulation. We performed atomistic simulations, totalling 0.5 milliseconds, on H, K, and NRas isoforms in the apo, GTP, and GDP-bound states of Ras protein, with and without complexing to its downstream effector, Raf. We show that differences in experimentally measured intrinsic and Raf-dependent catalytic speed amongst the three isoforms can be explained by dynamical activity and entropy, respectively. We show that Switch I and Switch II are the primary components of thermodynamic and kinetic allosteric networks, consistent with the key roles of these two motifs. These communication networks are altered by the hydrolysis of the substrate gamma phosphate, leading to increased entropy in HRas loops involved in substrate release. We find that the putative allosteric region is not coupled in KRas, but is coupled to the hydrolysis arm switch II in NRas and HRas, and that the mechanism in the latter two isoforms are thermodynamic and kinetic, respectively. Binding of Raf-RBD further activates thermodynamic allostery in HRas and KRas but has limited effect on NRas. These results indicate that kinetic and thermodynamic correlations are both needed to explain protein function and allostery. These two distinct channels of allosteric regulation, and their combinatorial variability, may explain how subtle mutational differences can lead to diverse regulatory profiles among enzymatic proteins.

## I. Introduction

Protein can transfer motion, and thus information, across the length of their structures. However, a mechanistic understanding of how this information flows is not yet established. While almost all drugs target protein active sites [1], very few target allosteric sites. With greater understanding of protein communication networks, new allosteric targeting sites could emerge for proteins that have eluded current drug screening methods. On a more fundamental level, a mechanistic understanding of allostery is required to build testable models of protein function and evolution. The first models for allosteric communication were phenomenological. The MWC model [2], posited that ligand binding shifted the equilibrium between an active and an inactive state, and the KNF model [3] posited that the protein’s flexibility allows the ligand to bind with an ‘induced fit’. In 1970, experiments by Perutz [4] were the first to use crystal structures to decipher allosteric mechanisms based on structural changes in hemoglobin upon binding of oxygen to the heme groups. Allostery was initially described in proteins with well-defined backbone structural changes between the active and inactive states. However, many researchers have since observed allosterically induced functional change without a difference in mean structure between functional states. For example, in the catabolite activator protein (CAP), two ligand binding sites communicate allosterically to change the protein’s affinity for DNA. This communication has been shown by NMR to be purely entropic [5]. As a complementary method, statistical analysis of pairs of amino acids that were jointly mutated in multiple sequence alignments were used to reveal contiguous sectors of coevolving residues within proteins [6, 7]. Through clever attachment of a light-activated protein domain at surface sites of the sector in DHFR protein, allosteric communication through the sector was demonstrated [8]. These results indicate that allosteric mechanisms are more diverse and nuanced than what can be inferred from static structure.

Our current understanding of allostery has evolved from a two-state model (i.e. the protein is either in the active state, or the inactive state) to a statistical ensemble-based model in which a protein samples a probability distribution over conformational states. Characterizing conformational states instead of relying on a single averaged protein structure, such as is done with x-ray crystallography, has proven to be important for understanding reaction mechanisms [9, 10]. Working under this conformational ensemble model, Cooper and Dryden mathematically showed that allostery can occur without enthalpic changes.[11] More specifically, they demonstrated two other possibilities: (i) allosteric regulation can modulate the number of conformational states accessed (the entropy), or (ii) allosteric regulation can modulate the waiting time between conformational changes. The first case (entropic) is sometimes referred to as “dynamic allostery”, although we will denote this as “entropic allostery” and use it interchangeably with the more general term “thermodynamic allostery”; we will sometimes call the mutual information (entropy) “thermodynamic correlations.” We will refer to the second possibility proposed by Cooper and Dryden as “kinetic allostery.” Entropic allostery has been corroborated by structural Nuclear Magnetic Resonance (NMR) and kinetic ITC studies [12,13,14] showing that conformational entropy can have a greater effect on binding affinity than differences in backbone structure. This well-supported abstract framework underscores the need to understand and predict networks of thermodynamic and kinetic correlations from structural and dynamical data on real systems.

Molecular Dynamics (MD) simulations provide an atomic level resolution of protein motions on the functionally relevant timescale of pico to milliseconds. The method is uniquely able to provide correlated motion of all protein atoms, which is need to predict allosteric networks even in the absence of allosteric ligand. In equilibrium MD, these correlations are sampled by collecting many instances of reversible stochastic fluctuations. Although some ligands can cause an “induced fit” change in protein conformation that differs from equilibrium protein fluctuations, there is also evidence that proteins populate alternate conformational states determined by intraprotein correlations, and that ligand binding “conformationally selects” for one of the states suited to a particular function. Thermodynamic couplings within the protein can be quantified by calculating the cross-correlation or covariance between the cartesian coordinates [15] and often further simplified by projecting them onto lower-dimensional principal components [16]. Because dihedral coordinates represent protein degrees of freedom subject to covalent bond constraints, they are the natural internal coordinates to represent protein conformations [17], and correlations between their non-monotonic probability distributions are well-represented by calculating the mutual information between dihedral coordinates [18, 19]. Pathways of information transfer have also been inferred using graph theoretic approaches such as the edge-betweenness measure to identify possible information bottlenecks connecting different protein regions [20]. However, relatively little progress has been made to quantify kinetic allostery, which would require capturing correlations in the changes of timescale of motion rather than the motion itself.

### Kinetic vs thermodynamic allostery

Because entropy and enthalpy are time-independent equilibrium measures, allosteric mechanisms driven by enthalpic and entropic changes can be quantified by probing the protein ensemble without considering the arrow of time. However, a kinetic measure must depend on time. Furthermore, to quantify kinetic allostery requires discrete event statistics in order to measure correlations between the frequency of conformational dynamics. Although methods exist to augment thermodynamic correlations with temporal information [19], we previously introduced a purely temporal measure of correlations in the waiting time between protein dihedral changes, which are well approximated as discrete jumps between distinct configurations [21]. We used this measure to demonstrate that side-chain-side-chain kinetic allostery is long-range (up to the protein length), whereas side-chain-side-chain thermodynamic allostery is locally confined [21]. Here, we apply this framework to half a millisecond of atomistic MD data to quantify backbone kinetic allostery and compare this to the thermodynamic allosteric network. Using the three main isoforms of the Ras family as model systems, we systematically studied the effect of ligand state and downstream effector binding on kinetic and thermodynamic correlation networks. We show that both are needed to explain differences in intrinsic versus Raf-dependent catalysis rates amongst the isoforms, and both types of networks predict coupling of the switches to the allosteric site, although the different isoforms vary in their use of kinetic versus thermodynamic allostery.

## II. Quantifying kinetic allostery

Given a protein with an allosterically de-activated state A, which then transitions to an allosterically activated state B, with each type of allostery, a different change in the energy landscape of the conformational ensemble is expected (Fig 1a). Enthalpic allostery results from changing the relative stability between states (e.g. in hemoglobin [4]), whereas entropic allostery changes the number of conformations accessible to one of the states (e.g. tetracycline receptor [10]; Fig. 1a). Kinetic allostery results from raising or lowering the energy barrier between states, resulting in changes to the waiting time between subsequent inter-state transitions. Parts of the protein with long mean waiting times indicate regions of dynamical frustration; if the waiting times of different parts of the protein vary in time in a correlated manner, they constitute a kinetic correlation network. If this correlation network includes surface sights whose waiting times can be modulated by external pertubation, then such a correlation network is a kinetic allosteric network. The influence of thermodynamic sources of allostery, whether entropic or enthalpic, on functional states is relatively intuitive. However, thus far the influence of waiting times as a cause of functional change has not been explored. We demonstrate below how changes in dynamical frustration, and therefore kinetic allostery, is sufficient to change the function of an enzymatic protein.

**Figure 1:**
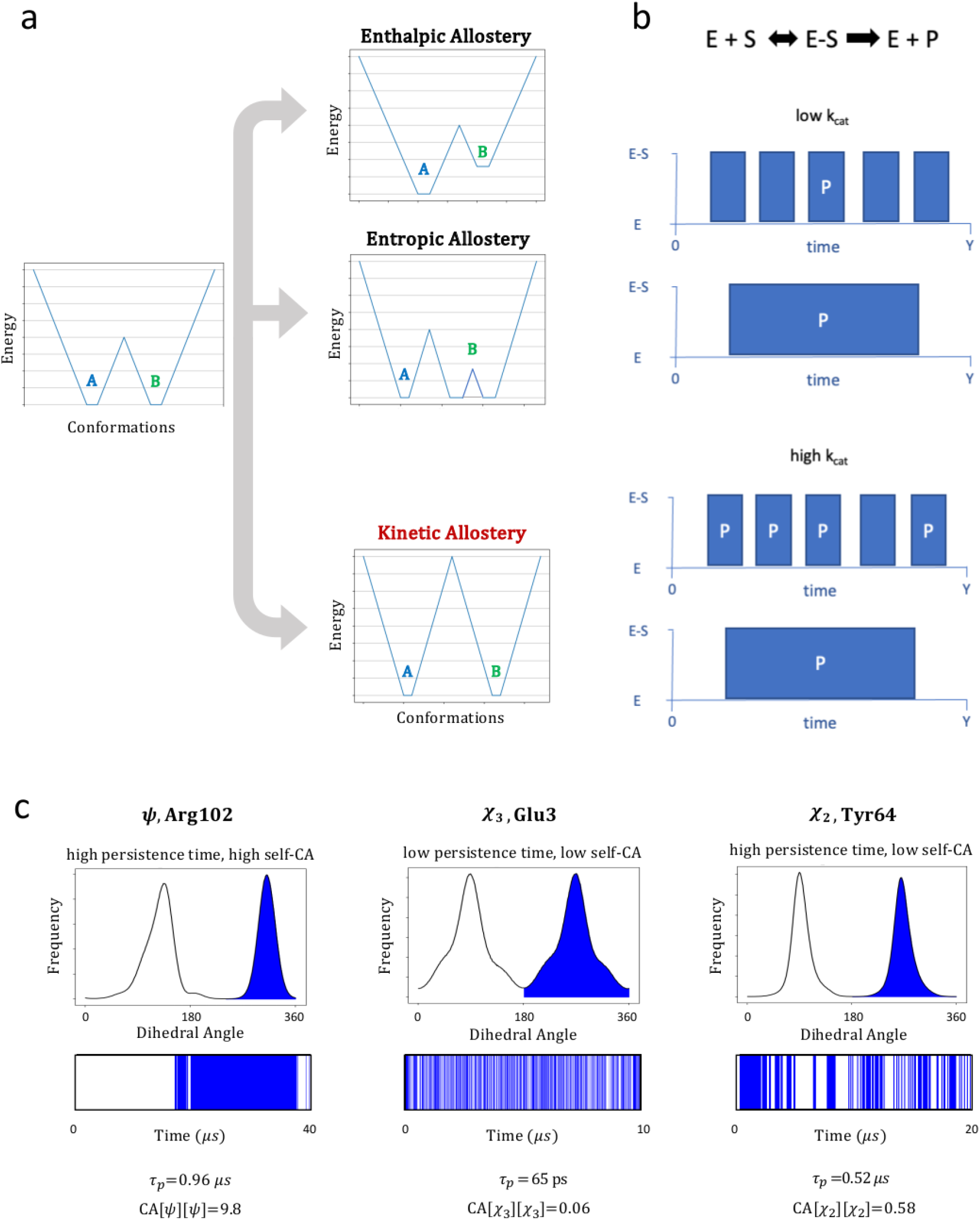
Types of Allostery. Timing Based Allostery, a. Energy landscapes for two protein states, A and B, where A is an inactive state and B is an allosterically activated state. Enthalpic and entropic allostery correspond to changing the relative energy or number of conformations (i.e. microstates) of state B, respectively. Kinetic allostery can be thought of as a change in the barrier height between states. b. Michaelis Menten kinetics (details in text) for two enzymes, one with a high k_r_ and k_f_, and one with low k_r_ and k_f_, but the same ratio of k_r_/k_f_. If k_cat_ is low, enzyme turnover rate is independent of k_f_. For high k_cat_, turnover rate becomes k_f_-dependent. c. Sample dihedral distributions from HRas MD simulations. For 3 very similar distributions, vastly different persistence times (τ_p_) and self-conditional activity (CA[x][x], i.e. dynamical memory) are present.

### The conditional activity (CA) as a metric for kinetic allostery

We do not expect waiting times (i.e. kinetic changes) to affect the activity of a protein that performs a reversible task that reaches local equilibrium, such as binding a target, because only the ratio of forward and backward reaction rates matters in that case. However, if the protein performs a non-equilibrium function, such as an irreversible enzymatic reaction (Figure 1b), altering waiting times can influence function. For example, in the standard Michaelis-Menten kinetics model of enzymatic reactions, the Michaelis constant *K_M_* is the concentration of substrate for which the enzyme achieves half of its maximum reaction rate (*V*_max_). *K_M_* is related to the microscopic kinetic rate constants [22]: *K_M_ = k_r_/k_f_* + *k_cat_/k_f_*, where *k_r_/k_f_* is the thermodynamic term that represents the probability that the enzyme is unbound, and *k_cat_/k_f_* is the kinetic term that represents the quotient of the rate of catalysis once the ligand is bound and the rate of binding. If the enzyme is coupled to a slow irreversible process (small *k_cat_*), the thermodynamic term *k_r_/k_f_* dominates, and *k_cat_* does not significantly influence the rate of formation of product. However, if the enzyme is coupled to a *fast* irreversible process, the kinetic term (*k_cat_/k_f_*) dominates. In the latter scenario, a change in waiting times that alters *k_f_* can influence the rate of product formation by influencing the likelihood that a binding event will result in product formation (Fig 1b). This is true even if the *k_r_/k_f_* ratio is unchanged. For such a catalytic protein, one would expect that long-range modulation of *k_f_* can serve as a purely kinetic form of allosteric regulation.

Kinetics of conformational transitions are the result of motions at several timescales: side-chain rotations and small loop motions over picoseconds to nanoseconds, and larger domain movements over microseconds to milliseconds [23]. The fastest functionally relevant protein motions are ps-ns motions, and have been shown to play a role in allostery for several systems [5, 24, 25, 26].

In contrast to well-studied measures of thermodynamic correlations, we quantify kinetic correlations using the conditional activity (CA) metric we previously introduced [21]:

The CA [21] is defined as

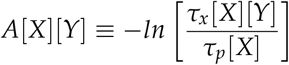

where (*τ_p_[X]*) is the persistence time and (*τ_x_ [X][Y]*) is the exchange time. The persistence time is (*τ_p_[X]*) the amount of time that passes, or the *waiting time*, between a random time point and a transition in X, and the exchange time is (*τ_x_[X] [Y]*) the waiting time between a transition in X and a transition in Y. In other words, given a time series of switching events of discrete variables *X* and *Y*, the conditional activity of *X* on *Y* is the log of the reduction of the expected waiting time between *Y* events immediately following an *X* event. Therefore, processes that are dynamically correlated and anti-correlated will have positive and negative CA, respectively (See methodological details in [21]).

Currently, most protein structural studies are based on Cartesian coordinates, where there is no threshold that can be used to determine if an event occurred. Calculating CA requires a reliable measurement of the timing of an event, and the degrees of freedom must reside in a finite set of discrete states. Protein dihedral angles, measured using microseconds-long MD simulations, meet this criteria because they sample probability distributions that are well-separated into distinct basins. The transition time of a dihedral conformation change is then the time in which it hops from one dihedral basin to another (see methods for vibrational filtering and transition commitment threshold). In prior work, sidechain CA was observed to have long-range correlations across the length of entire proteins, whereas sidechain mutual information was strictly short-range [21]. Here, we compare and contrast thermodynamic and kinetic allostery between backbone dihedral angles. It is important to note that we are using equilibrium sampling of each of the Ras states to discover correlated networks indicative of allosteric pathways controlling the non-equilibrium steps of the Ras cycle.

### Generating discrete conformational barcodes from MD simulations

To calculate the conditional activity measure of kinetic allostery, statistics of transition events are required, necessitating the construction of a ‘dihedral barcode’ for each time point, in which every position on the barcode represents a different backbone dihedral, and the discrete value of that position corresponds to the dihedral’s basin at that time point. The barcodes were constructed from the MD simulation data by first extracting dihedral angle probability distributions over the MD trajectory (Fig.2a, top). Dihedral peaks were assigned using a Matlab peak-finding function [27], with each peak corresponding to a discrete dihedral state (Fig. 2b), and assignment of a dihedral value to a dihedral state were based on the peak closest to the dihedral value, generating a discrete conformational barcode over time (Fib. 2a, bottom).

**Figure 2:**
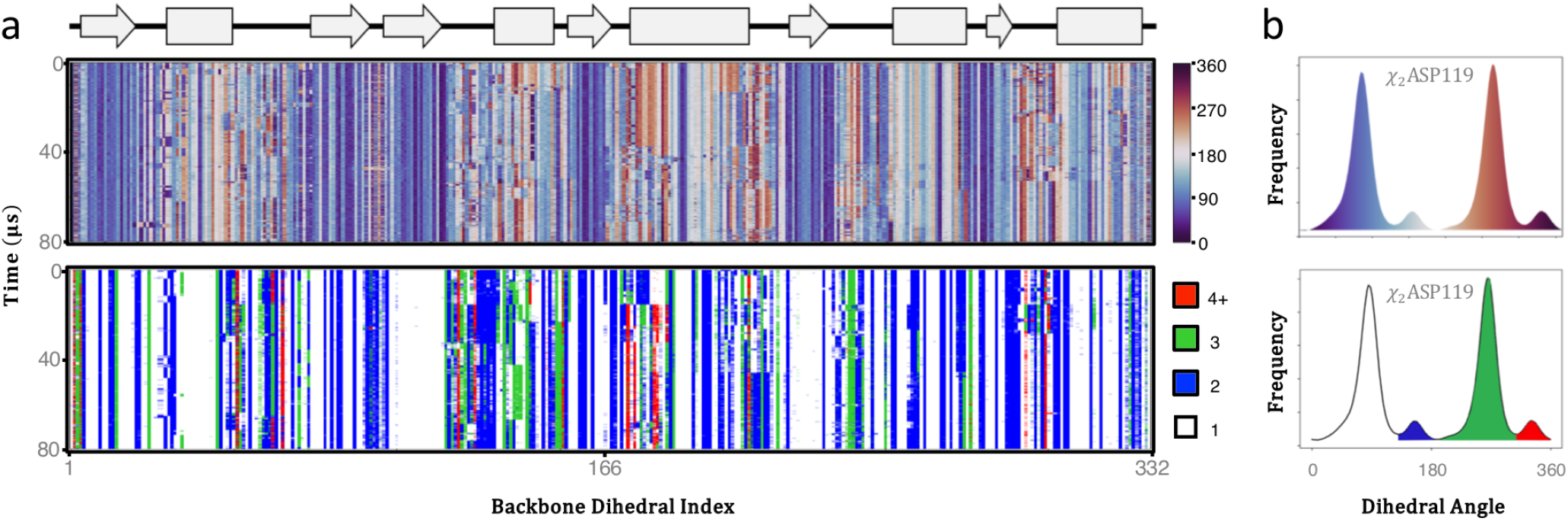
Types of Allostery. Vibrational filtering and barcoding a. HRas 80 microsecond GROMOS simulation data. Conversion of vibrational backbone dihedral data (top) into dihedral data that is binned by basin index(bottom). Each dihedral basin is assigned a number from 1-N, where N is the total number of basins observed for that dihedral distribution. Here each basin from 1-N is assigned a different color, unless there are more than 4 basins, in which case the 4th and greater basins are all red. The 166 amino acid protein has 332 backbone dihedrals (phi+psi for each amino acid) and are shown here in order of protein sequence. Secondary structure map at top with beta strands (arrow), alpha helices (rectangle) and loops (line). b. Example of vibrationally unfiltered dihedrals (top) and vibrationally filtered and binned rotamers (bottom) from a single dihedral distribution.

## III. Results

### Ras protein family as a model system for kinetic allostery

Ras is a GTPase upstream of the MAPK/ERK pathway that regulates cell fate and proliferation in response to growth factor stimulation at the cell membrane. They are enzymatic proteins that go through a cycle of binding GTP (the active state), hydrolyzing it to GDP (the inactive state), and releasing the GDP into the cytosol. In the active GTP-bound state, the Ras switch I domain can bind to a Raf kinase (Fig. 3a), dimerizing Raf and thus activating its ability to phosphorylate MEK [28]. Following product release, the Ras active site is subsequently occupied by GTP due to tenfold GTP/GDP concentration in the cell [29]. Before binding to Raf, Guanine Nucleotide Exchange Factors (GEFs) accelerate the rate at which GDP is released from Ras and thus serve to activate Ras. GTPase activating proteins (GAPs) increase the rate of GTP hydrolysis to GDP, thus deactivating the protein. Ras uses switch I (residues 30-40) and switch II (residues 60-76) for GTP hydrolysis (Fig. 3a) as well as for binding of these effector proteins. Because the hydrolysis step of the Ras cycle is an irreversible non-equilibrium process, it is a viable candidate for kinetic allostery.

**Figure 3:**
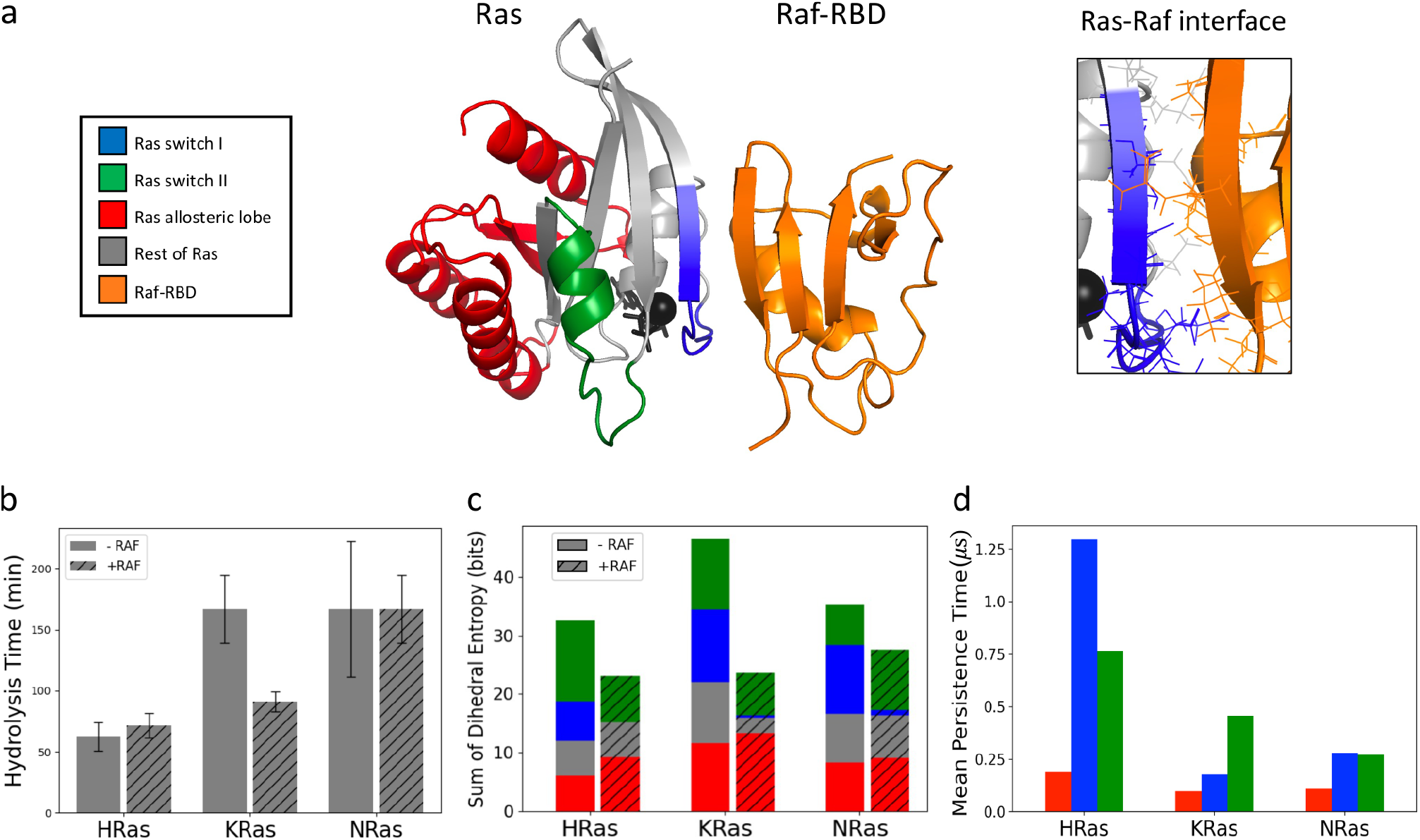
a. HRas-Raf RBD structure (PDB ID:4G0N). Allosteric lobe is represented in red, switch I, switch II in blue and green respectively. Raf-RBD is represented in orange. View of Ras-Raf-RBD interface at switch I shown on right b. Experimental hydrolysis time data adapted from Ref. [31]. Sum of individual dihedral entropies of Ras showing contribution from each domain of the protein d. Mean persistence time for each domain of Ras. This Ras-Raf structure shows the RBD. THere are two domains in Raf that bind to Ras: RBD and the CRD. The CRD was disordered in the crystal structure and is thus omitted. The CRD binds on either side of the RBD binding site of switch I.

Ras has been studied in depth because it is one of the most commonly mutated proteins across cancer [30]. Nevertheless, fundamental properties of Ras function and regulation are not mechanistically understood. There are three Ras isoforms: HRas, KRas, and NRas. The catalytic lobe of the protein (residues 1 through 86), which contains the hydrolysis machinery, is 100% conserved between isoforms, while the allosteric lobe of the protein (residues 87-166) is 80% conserved between isoforms. Despite this conserved catalytic lobe, the isoforms exhibit different hydrolysis rates [31]. Ras isoforms have different mutation rates and associated cancer types, localize to different membrane microdomains and subcellular compartments, control different transcriptional networks, and are modulated with different sensitivities by GAPs and GEFs [32, 33]. These differences in regulation suggest that there are distinct communication networks in the three isoforms despite sequence similarity. Ras proteins have been difficult to target pharmacologically due to the dearth of observed pockets on the protein surface, and the mechanism of Ras dimerization and activation of Raf is still not well understood. Nevertheless, a putative HRas allosteric site was identified at the N terminus of switch II, which is suspected to function in the Raf-bound state [34, 35]. While the endogenous ligand is not known for this site, it is suspected to play a role in Ras function based on observed binding of calcium and acetate, which results in an ordering of switch II that places Q61, the main amino acid assisting in hydrolysis, into the active site [34, 35]. This allosteric site is suspected to modulate hydrolysis in the presence of Raf because binding at this site results in a conformational change in switch II, which is outside the Raf binding site. The Ras family proteins are therefore ideal model systems to compare kinetic versus thermodynamic allostery, and to use these networks to explain differences in protein function arising from mutational variation, as well as differential regulation of protein function by ligand and cofactor binding.

### Ras molecular dynamics simulations

All-atom MD simulations were performed for Ras isoforms in their apo, GTD-bound, and GTP-bound states; the GTP-bound states were further divided into simulations with and without binding to Raf-RBD (See Table 1 and Methods). There are two splicing variants of KRas, KRas4A and 4B, and here we simulate KRas4B. The initial structures for the simulations were obtained from the PDB(https://www.rcsb.org) (See Methods section for system preparation and simulation details).

**Table 1.** Protein systems simulated. Slowly hydrolyzing GTP analogue GNP from PDB files were replaced with GTP prior to simulation. Missing atoms of Raf-RBD were also modeled.

| System | PDB | Simulation Time (μs) |
| --- | --- | --- |
| H-Ras Apo | 5P21 | 53 |
| H-Ras GTP | 5P21 | 60 |
| H-Ras GDP | 1CRQ | 51 |
| K-Ras Apo | 6GOD | 35 |
| K-Ras GTP | 6GOD | 50 |
| K-Ras GDP | 6MBT | 52 |
| N-Ras Apo | 5UHV | 47 |
| N-Ras GTP | 5UHV | 47 |
| N-Ras GDP | 5UHV | 41 |
| H-Ras GTP + Raf RBD | 4G0N | 16 |
| K-Ras GTP + Raf-RBD | 6GOD | 17 |
| N-Ras GTP + Raf-RBD | 5UHV | 17 |

### Correlating Raf-dependent simulation dynamics to experimental catalytic rates across Ras isoforms

Raf is a downstream effector of Ras that binds near the active site of Ras at switch I (Fig. 3a), and only binds when GTP is bound [36]. Experimental measurements of the GTP hydrolysis times of the three isoforms [31], both bound and unbound to Raf, are shown in Fig. 3b. These kinetic measurements reveal two trends among Ras isoforms. The first is that in the absence of Raf, HRas hydrolyzes GTP significantly faster than KRas and NRas. The second is that, while KRas hydrolysis speed is significantly faster when bound to Raf, binding of Raf does not affect hydrolysis rate of HRas and NRas.

We simulated wild-type GTP-bound NRas, KRas, and HRas, both bound and unbound to Raf-RBD, and calculated dihedral angle entropies and waiting time statistics. We find that configurational entropy can partially account for the differences in intrinsic and Raf-bound hydrolysis rates across isoforms. The backbone dihedral entropy, summed across the protein, is highest in KRas (Fig. 3c). Binding of an effector protein is expected to increase the rigidity of the protein at the binding site, as fluctuating residues in the pre-bound state become locked into a single conformation to stabilize the interaction with the effector protein. Upon binding of Raf, for all isoforms, the summed dihedral entropy decreases (Fig. 3c) across all isoforms, with the largest decrease in KRas (Fig. 3c). This supports the previous hypothesis that the high fluidity of KRas requires Raf to stabilize switch I in order to stabilize the surrounding GTP hydrolysis machinery [31], and further shows that Raf binding significantly reduces switch II entropy despite no direct contact with Raf, indicating thermodynamic correlations between switch I and switch II (see mutual information analysis below).

Although the differences in intrinsic and Raf-dependent hydrolysis rates between HRas and KRas can be explained by trends in entropy, NRas entropy is very similar to that of HRas, and yet has half the hydrolysis rate (Fig. 3b,c). The differences in intrinsic hydrolysis rates amongst the isoforms, including the descrepancy between HRas and NRas, can be explained by the persistence times measured in the GTP-bound simulations (Fig. 3d). Persistence time is the mean waiting time until a dihedral transition–a switch from one dihedral basin to another–starting from a random time point. The persistence times within a structural motif such as switch I is a measure of dynamical frustration, which can lead to long residence times in configurations necessary for catalysis. HRas’ mean persistence times in the hydrolysis machinery (switches I and II) are an order of magnitude greater than that of NRas and KRas (Fig. 3d). Although NRas and HRas entropies are similar (Fig. 3c), the higher persistence time of HRas can account for the discrepancy between HRas and NRas hydrolysis rates (Fig. 3b), suggesting that HRas’ intrinsically fast catalysis is enabled by slowing down the fluctuations of its switches.

These data suggest that dihedral persistence times and entropy are both needed to explain hydrolysis rates in Ras, with the former determining intrinsic rate and the latter determining the Raf-dependent rate. These indicate that thermodynamic and kinetic analyses are both necessary to explain protein function, even within a single protein, and motivate the analysis of waiting time correlations between dihedrals to infer kinetic allostery.

### Raf-dependent allosteric pathways across Ras isoforms

For each of our simulations, we calculated the inter-dihedral conditional activity and mutual information matrices to determine kinetic and thermodynamic pathways of allostery, respectively. In all Ras isoforms and regardless of nucleotide binding, the MI and CA communication networks (Figs. 3, S1, and S2), switch I and switch II contain the greatest amount of correlation, and this communication is mainly inter- and intra-switch communication. The MI and CA matrices in Fig. 3 also demonstrate that the presence of the gamma phosphate substantially alters communication networks in HRas, and this phenomenon is consistent across isoforms (Figs. S1 and S2). The GDP and apo (Fig. S6) states both contain greater amounts of communication than the GTP state in the allosteric lobe, indicating that the gamma phosphate quenches some communication networks within the protein. In the GDP-bound state, the function of Ras is mainly to release the GDP into the cytosol so that a GTP can bind in its place. This release function involves the coordination of residues in the allosteric lobe, including the breaking of a salt bridge between loop 8 (red, Figure 3) and loop 10. These loops contain conserved [37, 38] NKCD (143-147) and ETSAK (116-119) motifs, which are also responsible for the specificity of the guanine nucleotide and stabilizing nucleotide binding, respectively. The increased communication observed in the allosteric lobe in the GDP-bound state matrices is consistent with this function of the GDP-bound state. Changing the ligand from GTP to GDP increases entropy of loops 7 and 8 in HRas (Fig. 4b) and loop 7 in KRas (Fig. S1b), with no significant change in NRas (Fig. S2b). This is consistent with the picture of loops 7 and 8 acting as entropic reservoirs that help to stabilize GDP-bound Ras. In HRas (Fig. 4b), loop 8 shares significant mutual information with switch I and switch II, suggesting that regulation of switch I and switch II conformations by GEFs could allosterically lower the entropic favorability of the GDP-bound state via loop 8. Orthogonal to this thermodynamic picture, kinetic correlations in HRas are oppositely affected by the change from GTP to GDP binding. Despite loop 7 having lower entropy and lower mutual information with the switches in the GTP-bound state (Fig. 4b, top), analysis of the conditional activity shows that the fluctuations in loop 7 are much more kinetically coupled to the fluctuations in the switches in the GTP bound state (Fig. 4b, bottom). In contrast, loop 7 is thermodynamically uncoupled to the rest of the protein in the GTP and GDP bound states (Fig. 4b, top). This suggests that allosteric modulation of loop 7 in the GTP-bound state in the absence of GAP binding is kinetically controlled in HRas. These thermodynamic and kinetic correlations are not observed in the other two Ras isoforms (Fig. S1 and S2), indicating that although trends in entropy are consistent across isoforms, mutual entropy and dynamical activity are more sensitive to mutational modulation.

**Figure 4:**
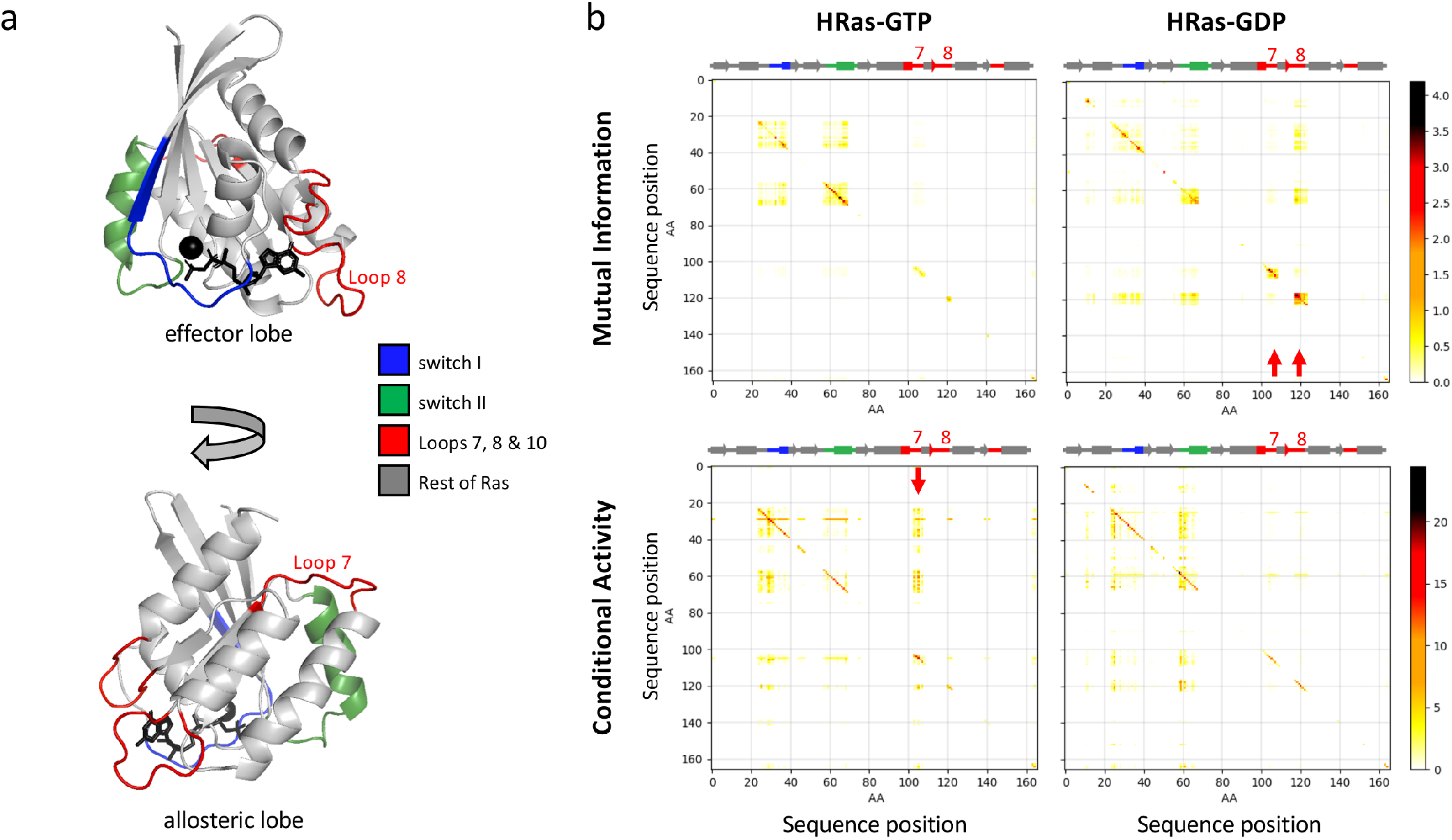
a. Two views of HRas (5P21) structure with important structural motifs labeled: switch I in blue, switch II in green, and loops 7, 8, and 10 in red. b. Mutual information and conditional activity for simulations of HRas bound to GTP and GDP. The corresponding NRas and KRas heatmaps are given in supplementary. Conditional Activities shown here are absolute values. Red arrows highlight differences in MI and CA for loops 7 and 8 upon GTP hydrolysis. Note different color scales for MI and CA.

For each of the Ras simulations, we diagonalized the MI and (symmetrized) CA matrices in order to find the principal eigenvectors. These eigenvectors show the most prominent CA and MI correlation networks for each simulation. For GTP-bound HRas, the magnitude of the principal eigenvector components of the *φ* and *ψ* dihedrals are shown proportional to the radius of their correponding C-alpha and carbonyl carbon atoms, respectively (Fig. 5a), and the corresponding eigenvalue compared to the other top eigenvalues of the spectrum (Fig. 5b). The main signaling function of Ras is to activate the downstream effector Raf. In the GTP-bound state, Ras can either bind to Raf (at switch I) or hydrolyze the GTP to GDP (involving switch I and switch II). Consistent with this expectation, the top MI and CA eigenvectors are dominated by switch I and switch II at the GTP-bound step of the Ras cycle (Fig. 5a, left). When the structure is simulated with Raf bound at switch I, the switch I dihedrals no longer participates in the dominant correlated network (blue spheres in Fig. 5a). These Raf-induced changes are similar for thermodynamic and kinetic networks at the level of the switches, despite variation in the specific dihedrals within the switches that contribute most to the networks.

**Figure 5:**
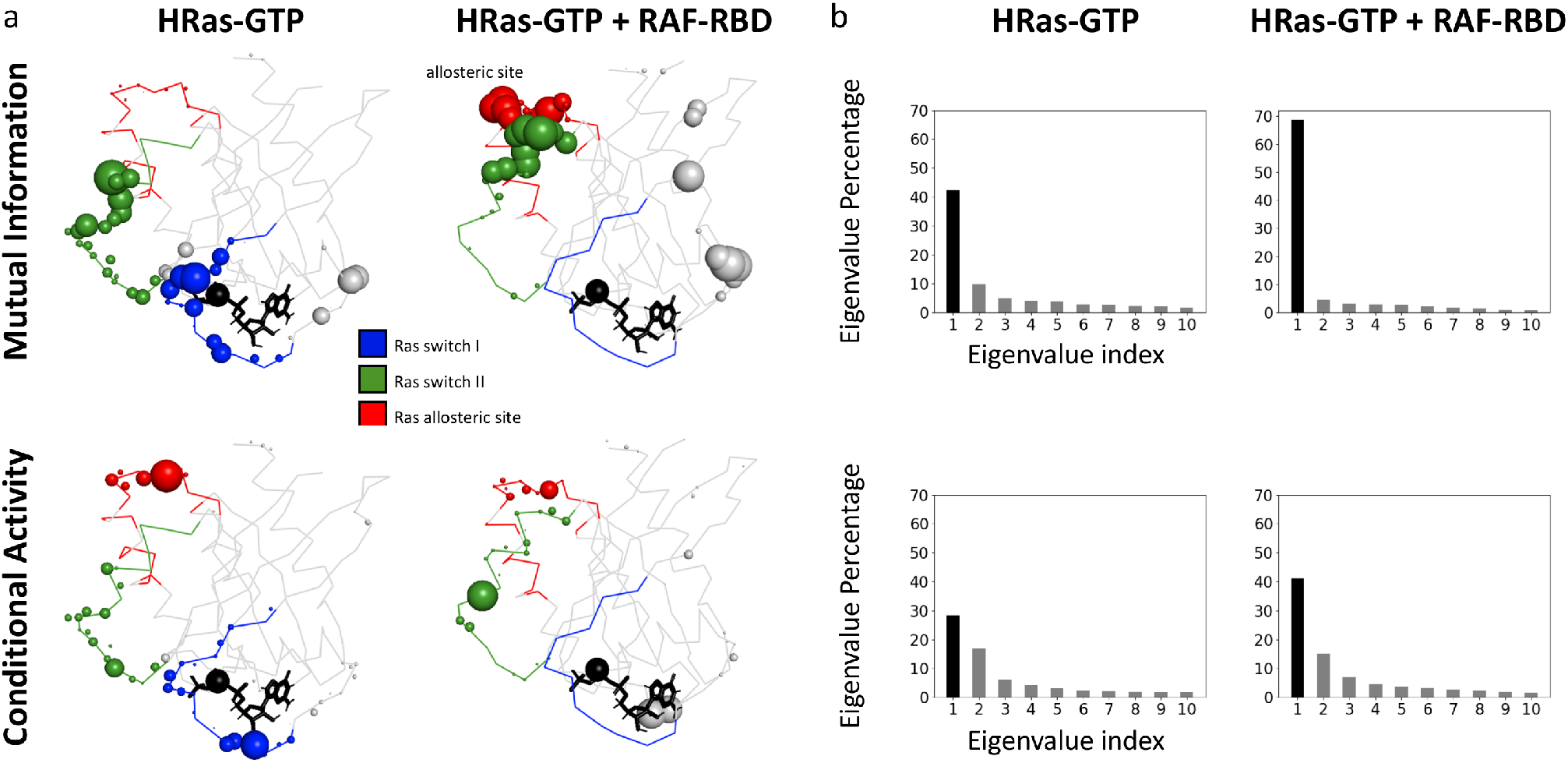
HRas Principal Eigenvectors. **a.** HRas-GTP and HRas-GTP +RAF-RBD MI and CA Principal Eigenvectors projected onto PDB structure (5P21). Sphere size is scaled to the size of the contribution of the nearest dihedral in the principal eigenvector. We use a minimum sphere size of 0.2 (CA or MI=0) and a maximum sphere size of 1.2, in order to represent all dihedrals visually, φ dihedrals are represented by C-alpha atoms and ψ dihedrals are represented by carbonyl carbon atoms. **b.** Top 10 Eigenvectors and their percentages of the eigenvalue.

In contrast, kinetic and thermodynamic communication networks, and their regulation by Raf, qualitatively differ in terms of the involvement of the allosteric site within loop 7 (red in Fig. 5a). Hydrolysis is suspected to be mediated by binding of a ligand at the allosteric site at the C terminus of switch II that functions exclusively in the Raf-bound state by ordering switch II and positioning Q61 in the active site [31]. In the absence of Raf, the principle eigenvector contains no intrinsic thermodynamic coupling to the allosteric site (Fig. 5a, top left), but significant kinetic coupling (Fig. 5a, bottom left) between the putative allosteric site and switch II. However, the Raf-bound simulations show that thermodynamic and kinetic allostery can be differentially activated by Raf (GAP) binding. Although Raf binding does not enhance kinetic allostery (Fig. 5a, right bottom), it is sufficient to activate thermodynamic allostery (Fig. 5a, right top). This differential regulation of kinetic versus thermodynamic allostery by Raf binding is also observed in KRas and NRas, but with isoform-dependent variation (Figs. S3 and S4). In particular, Raf binding activates both thermodynamic and kinetic allostery in KRas (Fig. S3), whereas Raf binding does not significantly affect allosteric engagement in NRas (Fig. S4).

## IV. Discussion

Cooper and Dryden were the first to demonstrate that kinetic changes, in addition to thermodynamic changes, can in principle tune protein function. However, in the almost 40 years since, how kinetic changes are coupled at the molecular scale to form allosteric networks remains largely unexplored. Computational advances increasingly allow interrogation of correlated motions responsible for allostery. In this study, we addressed this question by applying a recently developed measure of kinetic correlation to transition dynamics between dihedral states collected from atomistic simulations of the Ras protein family totally half a millisecond. In the case of the hydrolysis machinery, these networks are consistent across functional states of the protein (GTP-bound vs. GDP-bound) as well as across isoforms. The coupling analysis correctly identifies the important switches of the protein and their communcation with each other, as well as a previously suspected allosteric region. The analysis also reveals new insights into the nature of kinetic versus thermodynamic coupling between these region as well as key loop regions, and how these couplings differ amongst the Ras isoforms. Therefore, networks have both robust as well as tunable features. By comparing the resultant kinetic correlation networks with the mutual-information-based thermodynamic correlation networks, we demonstrate that kinetic control and kinetic allosteric networks are distinct from their thermodynamic counterparts. In fact, kinetic and thermodynamic characterizations are both needed in order to explain the diversity of cofactor-dependent catalytic activities and allosteric responses observed experimentally. The kinetic communication hotspots do not follow the boundaries of secondary structural elements, supporting the idea that dynamical modules are not necessarily structural modules [39].

This kinetic analysis framework opens up important new avenues of research in the study of allostery. Future work will include creating an online tool to make conditional activity analysis easily accessible. As computational capacity increases and molecular dynamics force fields become more reliable over longer simulation times, longer simulations will provide more insight to kinetic changes taking place over longer time periods, which will open up opportunities for studying larger proteins. In the case of Ras, further insight can be gained with simulation of additional mutated Ras systems, and with longer simulations of the systems presented here. An exciting path forward will be found in analyzing kinetic and entropic communication networks between multi-protein systems. An interesting future step for the Ras project will be to calculate the conditional activity in the Raf-RBD protein while bound to Ras, especially given that Raf is the downstream effector of this oncogenic protein. We hope that this framework will broaden understanding of possible allosteric mechanisms, and allow for prediction of new allosteric sites.

## V. Materials & Methods

### 1. MD Simulations

All-atom MD simulations were performed for Ras isoforms in their apo, GTP-bound, GDP-bound states, as well as the state of being bound to both GTP and Raf-RBD. There are two splicing variants of KRas, KRas4A and KRas4B, and we simulated KRas4B (referred to as KRas in this work). The initial structures for the simulations were obtained from the PDB and were altered as described in Table 1. Because Ras hydrolyzes GTP, GNP (phosphoaminophosphonic acid guanylate ester), a non-hydrolyzable analog of GTP, is used in crystal structures of Ras. For our GTP-bound simulations we replaced GNP by GTP by substituting the nitrogen with an oxygen atom in the PDB file. Simulations were performed with GROMACS [40, 41] at the BIOHPC computing facility at UT Southwestern. V-sites [42] were used to freeze hydrogen vibrations in apo simulations, allowing for a 5 fs timestep for those simulations, and 2 fs timesteps were used without V-sites for all non-apo simulations. Coordinates were saved every 10 ps for each simulation except for apo HRas and KRas simulations, which were saved every 25 ps. All simulations were run with the CHARMM36 force field [43] and a cubic periodic box with 1 nanometer solvent buffer distance was used (see Table S1 for system sizes). The SPC/E water model [44] was used and Na+ atoms were added to create an overall charge neutrality. The systems were energy minimized, heated to 300 K over the course of 10 ns in the NVT using the Berendsen [45] thermostat and pre-equilibrated for 1000 ns in the NPT condition using Nose-Hoover thermostat[46, 47] and Parrinello-Raman barostat [48]. A Verlet cutoff scheme was used for non-bonding interactions, and Particle Mesh Ewald (PME)[49] was used for long-range electrostatics calculations with the periodic boundary conditions. Pymol [50] was used for molecular graphics [50].

### 2. Data Coarse-Graining

Dihedral angle trajectories were used to generate probability distributions of each backbone and sidechain dihedral angle over time. Each amino acid has three backbone dihedrals: phi, psi and omega. Only phi and psi dihedrals were analyzed, since omega dihedrals stay in one rotamer. The peaks in these probability distributions were assigned using a MATLAB peak-finding function [27], and assignment of individual rotamers to peak basins were based on the peak closest to the rotamer. From this information, the raw 0-to-360 degree dihedral trajectories are converted into “peak trajectories”, where vibration in the raw data is filtered out (Fig. 2). Because occupancy of dihedral wells was insensitive to temporal resolution above 10 ns, for downstream calculation of mutual information, the 10 ps resolution peak trajectory is made into a 10 ns resolution consensus trajectory by finding the peak consensus within 10 ns time chunks. In contrast, we observed short-lived dynamical fluctuations below this time resolution, and consequently used the 25 ps resolution peak trajectories for conditional activity analysis. For ease of visualization, we used the 10ns consensus trajectory to build the ‘dihedral barcodes’ which represents the discrete backbone dihedral configuration as a function of time (Fig. 2a). Unless otherwise noted, programming was done using Python3 [51].

### 3. Correlation Analysis

The mutual information[52] between two dihedrals X and Y is defined as

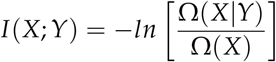

The conditional activity [21] is defined as

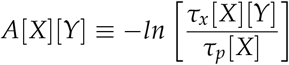

where *τ ≡ T*(*X, N(X)*) is the observation time. The transition time function is *T*(*X, i*), which is the time of the ith transition of *X*. The waiting time function is *W*(*X, t*), is the time interval, starting at time *t* until the next transition of *X*. The persistence time is 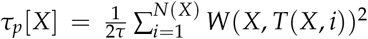 and the exchange time is 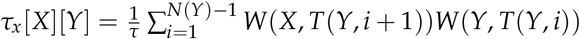.

## VI. Acknowledgements

The authors would like to thank Levent Sari for his help with MD simulations.

## VII. Supplementary information

**Table S1:**
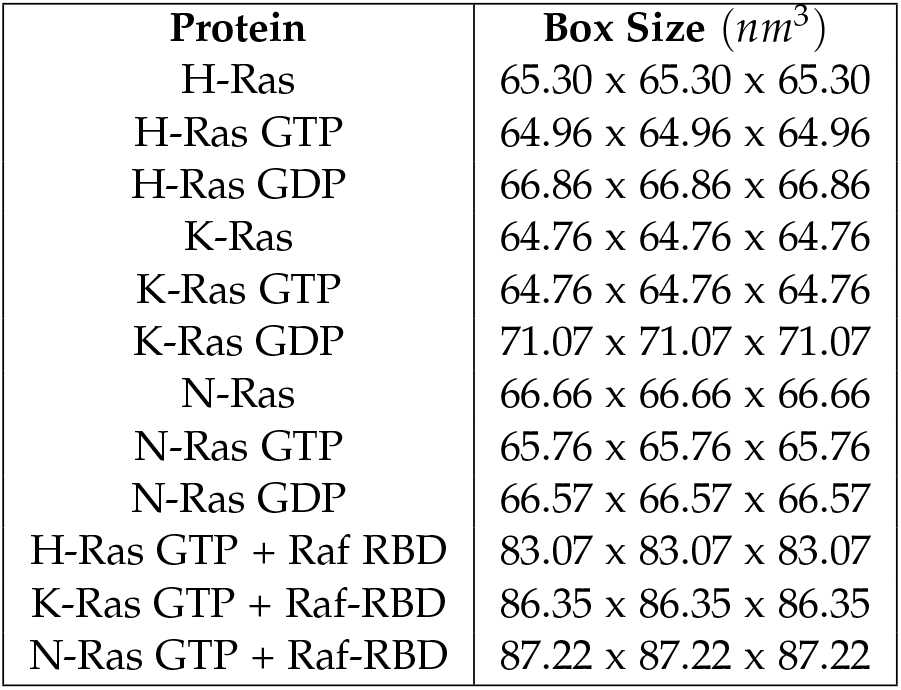
Protein simulation box sizes.

| **Protein** | **Box Size ($nm^3$)** |
| --- | --- |
| H-Ras | 65.30 x 65.30 x 65.30 |
| H-Ras GTP | 64.96 x 64.96 x 64.96 |
| H-Ras GDP | 66.86 x 66.86 x 66.86 |
| K-Ras | 64.76 x 64.76 x 64.76 |
| K-Ras GTP | 64.76 x 64.76 x 64.76 |
| K-Ras GDP | 71.07 x 71.07 x 71.07 |
| N-Ras | 66.66 x 66.66 x 66.66 |
| N-Ras GTP | 65.76 x 65.76 x 65.76 |
| N-Ras GDP | 66.57 x 66.57 x 66.57 |
| H-Ras GTP + Raf RBD | 83.07 x 83.07 x 83.07 |
| K-Ras GTP + Raf-RBD | 86.35 x 86.35 x 86.35 |
| N-Ras GTP + Raf-RBD | 87.22 x 87.22 x 87.22 |

**Figure S1:**
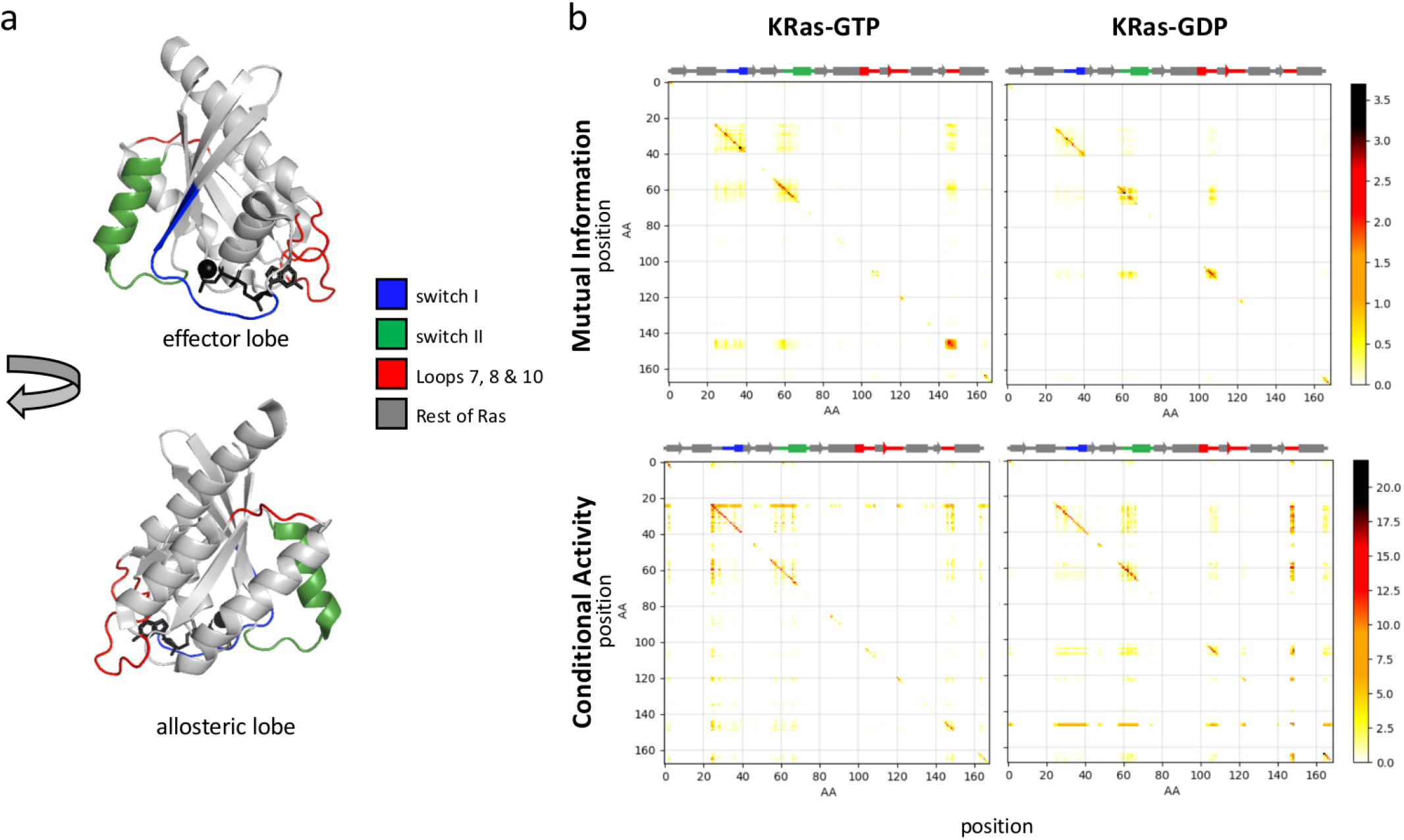
KRas structure bound to GTP (a). Mutual information and conditional activity matrices for KRas bound to GTP and GDP (b).

**Figure S2:**
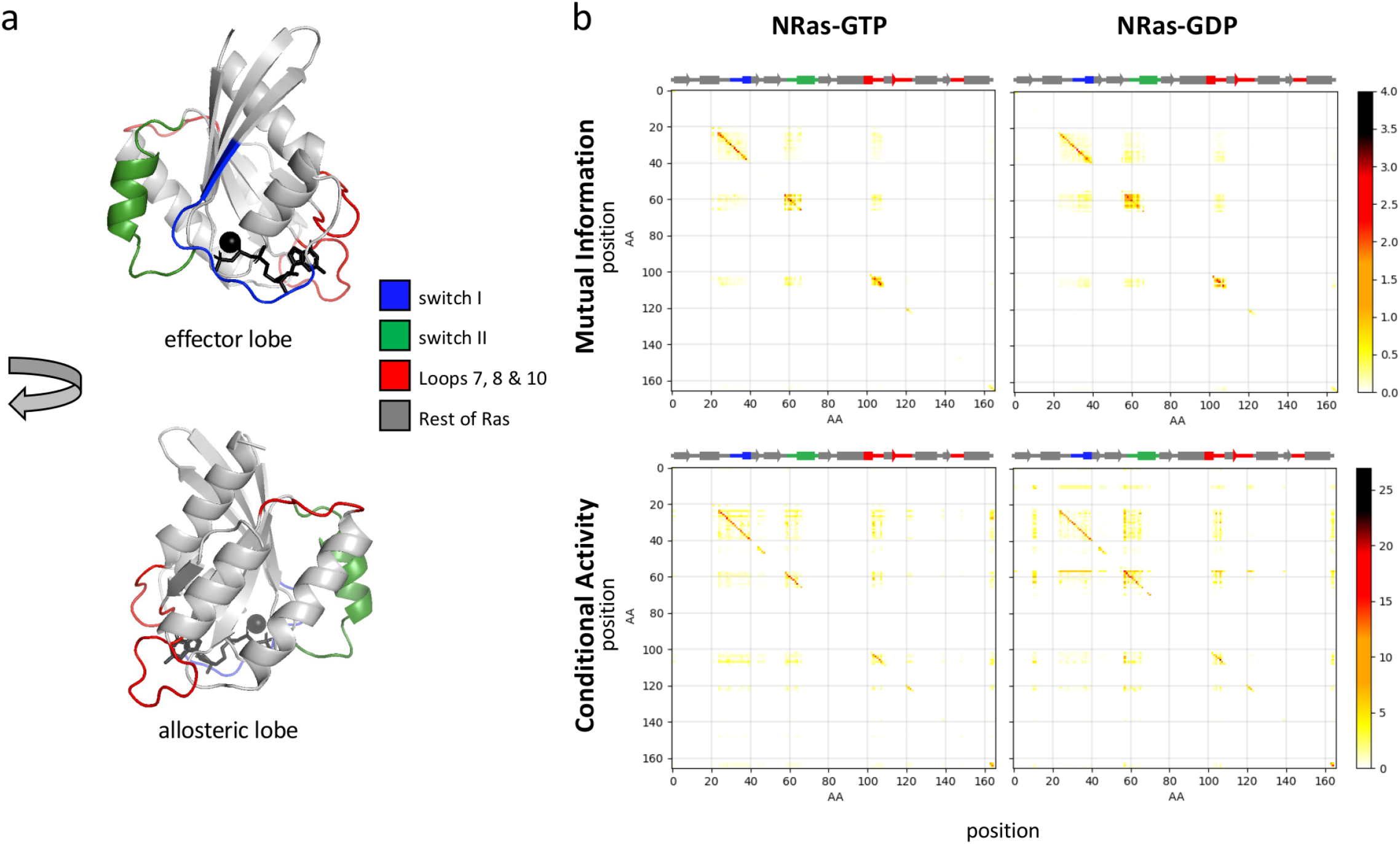
NRas structure bound to GTP (a). Mutual information and conditional activity matrices for NRas bound to GTP and GDP (b).

**Figure S3:**
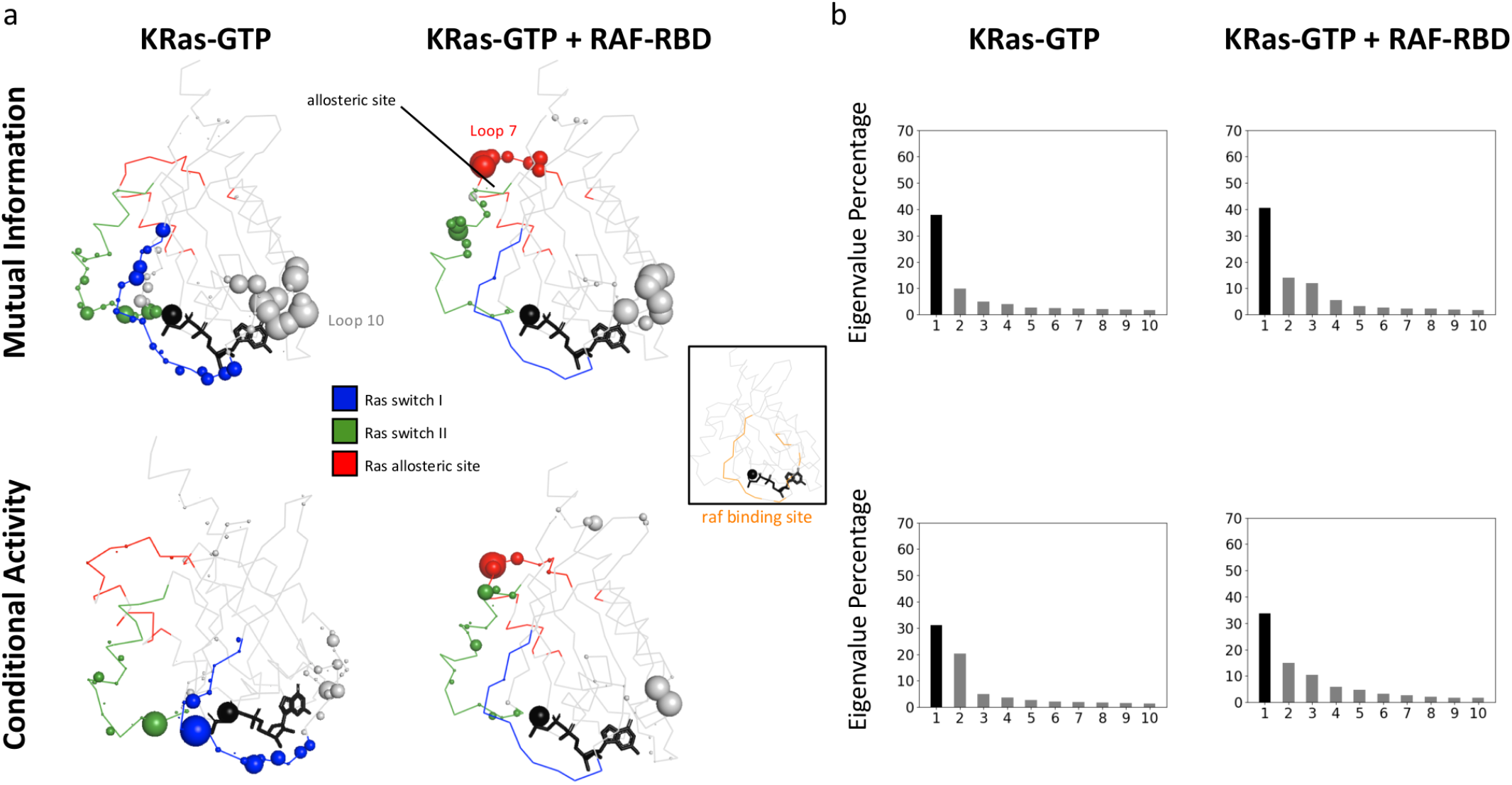
KRas-GTP principal eigenvector of the mutual information (top) and conditional activity matrices(bottom) are shown without (left) and with (right) Raf-RBD bound simulations (a). The eigenvalue fractions of the top ten eigenvectors are shown, with the principal eigenvalue corresponding to the eigenvectors shown in (a) colored black (b).

**Figure S4:**
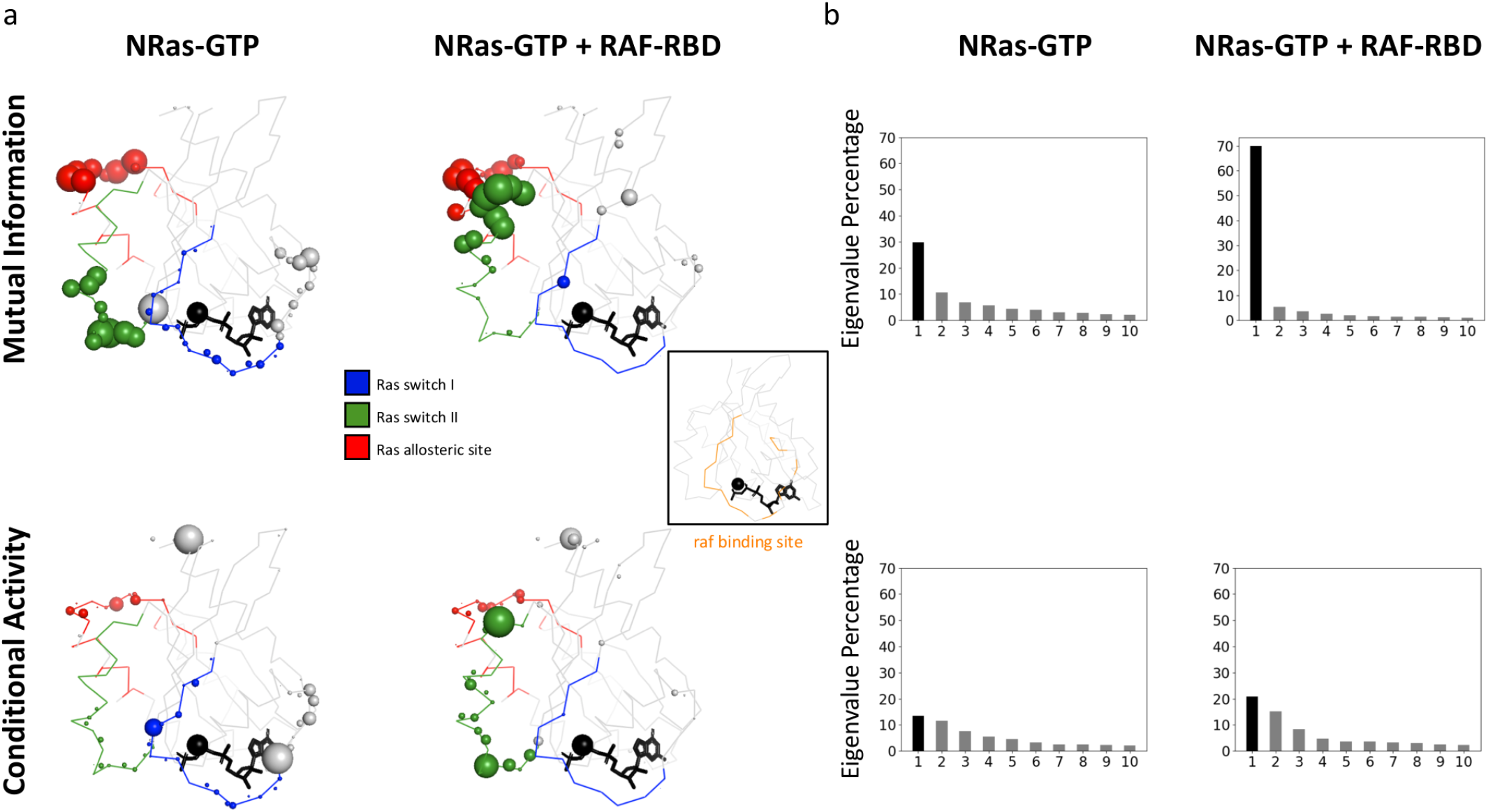
NRas-GTP principal eigenvector of the mutual information (top) and conditional activity matrices(bottom) are shown without (left) and with (right) Raf-RBD bound simulations (a). The eigenvalue fractions of the top ten eigenvectors are shown, with the principal eigenvalue corresponding to the eigenvectors shown in (a) colored black (b).

**Figure S5:**
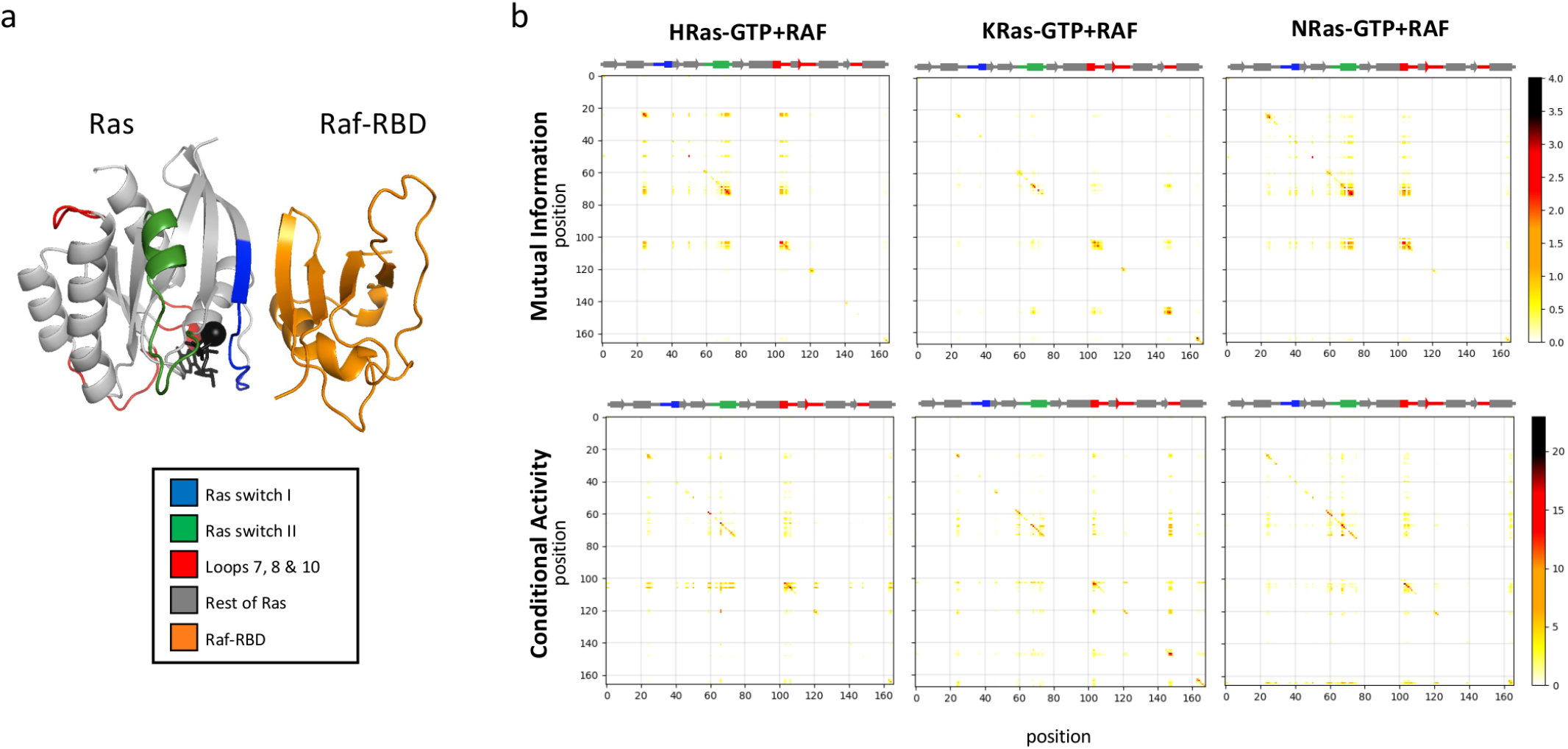
Ras-Raf complex (a). Mutual information and conditional activity matrices of HRas, KRas, and NRas bound to GTP and Raf (b)

**Figure S6:**
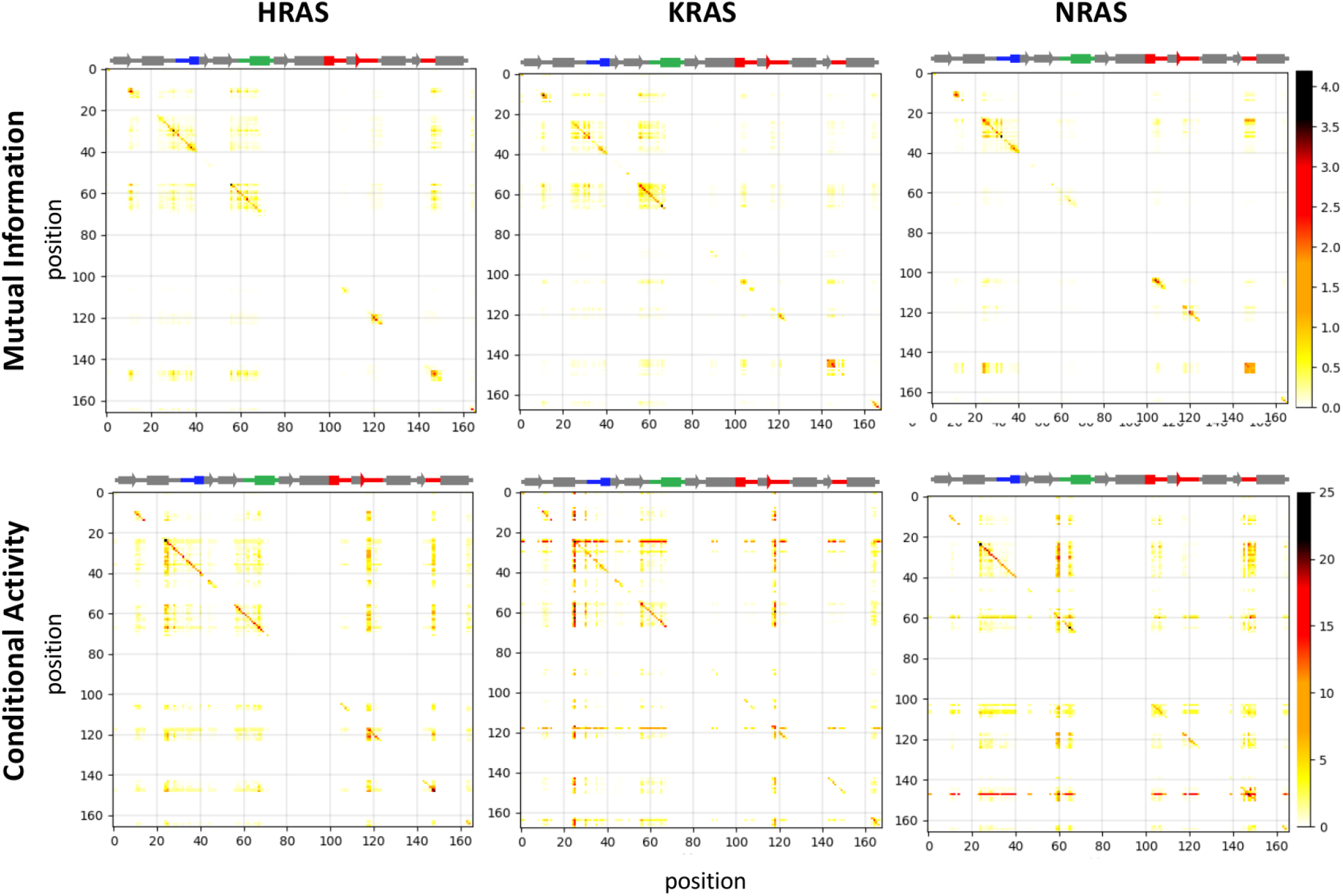
Mutual information and conditional activity matrices of HRas, KRas, and NRas in the apo state

